# A Rapid 10-Minute Silver Nitrate Staining Method for Visualizing the Osteocyte Lacuno-Canalicular System

**DOI:** 10.64898/2026.04.20.719546

**Authors:** Jinlian Wu, Libo Wang

## Abstract

The Ploton silver method employs a 50% silver nitrate solution (w/v; 2.943 mol/L) for staining and quantitative analysis of the osteocyte lacuno–canalicular system (LCS). We previously demonstrated that lower silver nitrate concentrations (0.5–1 mol/L) stain the LCS more effectively, revealing a greater number of LCS than the Ploton silver method. However, the staining duration of our initial modified method (60 minutes) remained comparable to that of the Ploton silver method (55 minutes), limiting its broader adoption. Here, we developed a rapid silver nitrate staining method by systematically evaluating the effects of temperature on staining efficacy. We found that incubation at 50–70°C for 10 minutes with a 1 mol/L silver nitrate solution produced optimal results. This rapid high–temperature method achieved excellent LCS visualization in bone samples from multiple animal species and in mouse pathological models. Moreover, high-temperature staining mitigated the LCS damage and insufficient staining associated with the 50% silver nitrate solution used in the Ploton silver method. This rapid 10-minute silver staining technique, designated the Wu-Wang silver method, provides a more accurate and efficient approach for LCS staining and quantitative analysis. Its adoption will facilitate systematic characterization of LCS morphological variations across vertebrate species, thereby advancing our understanding of osteocyte morphogenesis and the pathogenic mechanisms underlying bone and joint diseases.

**Graphical abstract** (Created in BioRender, https://BioRender.com)

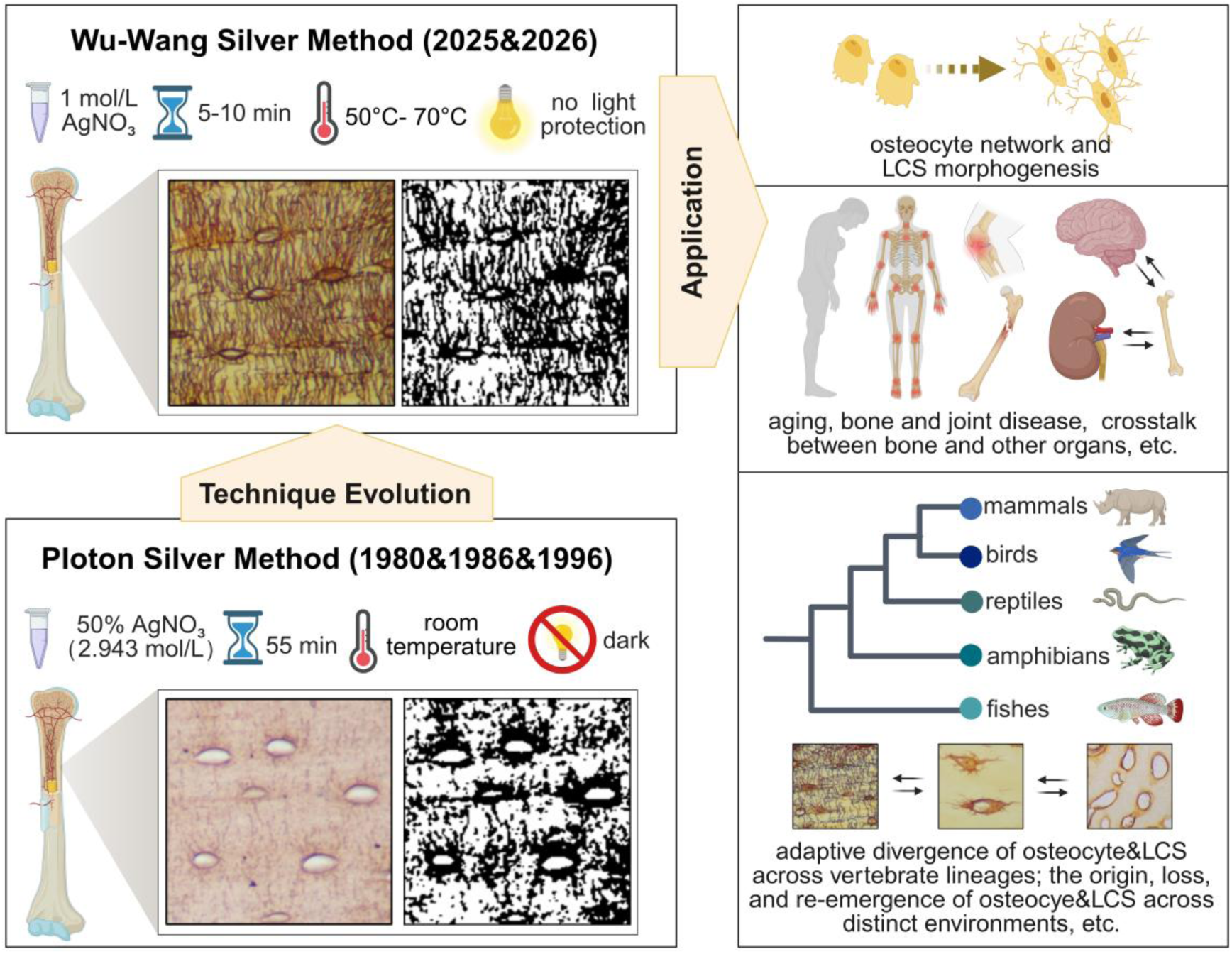

**Highlights:**

1. Elevating the staining temperature to 50–70°C enabled rapid and efficient silver nitrate staining of the osteocyte lacuna-canalicular system (LCS) within 5–10 minutes using 1 mol/L silver nitrate.
2. The high-temperature Wu-Wang silver method outperformed the conventional Ploton silver method, providing superior osteocyte LCS visualization while eliminating issues of osteocyte LCS damage and insufficient staining observed with the Ploton silver method.

## Introduction

In the mineralized extracellular matrix of vertebrate bones resides a cell type with striking morphological similarities to neurons: the osteocyte [1–2]. Osteocytes differentiate from osteoblasts and extend numerous dendritic processes from their surfaces [3]. These processes form an osteocyte network through intercellular coupling [4]. Analogous to electrical wires enclosed within protective conduits, this osteocyte network is intricately embedded and traverses within the lacunar–canalicular system (LCS)—an equally sophisticated architectural network [5]. The LCS consists of lacunae housing osteocyte cell bodies and canaliculi containing their dendritic processes and pericellular fluid. The structural integrity, connectivity, and permeability of the LCS critically influence osteocyte function and skeletal health [6–9].

Despite the pivotal roles of the osteocyte network and LCS in regulating bone remodeling and homeostasis, detailed morphological characterization and comprehensive understanding of LCS development remain challenging [10]. The Ploton silver method, initially developed for visualizing argyrophilic nucleolar organizing regions (AgNORs), was serendipitously adapted by Chappard *et al.* in 1996 for staining the osteocyte LCS and has since been applied in the field of osteocyte research [11–12]. However, its broader utility has been constrained by inconsistent staining performance and variability in reagent concentrations and compositions.

In our prior studies [13–15], we discovered that lower silver nitrate concentrations (0.5–1 mol/L), combined with optimized gelatin–formic acid solutions, achieved superior LCS visualization compared with the Ploton silver method (50% silver nitrate, w/v; 2.943 mol/L). We accordingly developed a novel optimized silver staining protocol with refined solution preparation and procedural steps [13–15]. Notably, we identified that the high silver nitrate concentration in the Ploton method resulted in osteocyte LCS damage and insufficient staining [13, 15]. However, given that the staining time of our optimized method (60 minutes) is comparable to that of the Ploton silver method (55 minutes), this may limit the acceptance and widespread application of our novel optimized method in the osteocyte biology community.

In the present study, we systematically investigated the effects of temperature on the staining efficacy of the LCS, building upon preliminary observations from our previous work [15]. We unexpectedly found that increasing the incubation temperature from room temperature to 50–70°C enabled rapid and complete LCS visualization within 5–10 minutes. Notably, these elevated-temperature conditions eliminated the issues of osteocyte LCS damage and insufficient staining associated with the 50% silver nitrate solution utilized in the Ploton method, as identified in our previous studies [13, 15]. We further demonstrated the applicability of this rapid method across diverse vertebrate species and in a murine model of impaired bone growth. The rapid method was validated in bone specimens of varying ages, subjected to different decalcification regimens, and treated with gelatin from different commercial sources and types. Collectively, these findings represent the first report of a rapid, efficient, and reproducible silver staining method for comprehensive LCS visualization. We propose that adoption of this method will substantially advance understanding of osteocyte morphogenesis, LCS development, and their relevance to skeletal pathology. Moreover, this approach provides a practical histological tool for systematically characterizing the presence or absence of osteocytes (cellular vs. acellular bone) and their LCS abundance (high vs. low abundance LCS bone) across different taxa (terrestrial vs. aquatic vertebrates, and advanced vs. basal teleosts) [16–18], and for investigating the evolutionary origin, loss, and re-emergence of these beautiful structures [19–20].

## Materials and Methods

### Reagents

0.5 mol/L EDTA decalcification solution (pH7.2, G1105-500ML, Servicebio, Wuhan), 10% EDTA decalcification solution(diluted from 0.5 mol/L EDTA), 14% EDTA decalcification solution(diluted from 0.5 mol/L EDTA), 5% Formic acid decalcification solution (G1107-500ML, Servicebio, Wuhan), 10% Neutral formalin fixative(311010014, Wexis, Guangzhou), 1 mol/L Silver nitrate solution(P1929026, Bolinda Technology, Shenzhen), Silver nitrate power (C510027-0010, Sangon Biotech, Shanghai), Type-B Galetin (G8061, bloom ∼225 g, Solarbio, Beijing), Type-A Galetin (A609764-0100, Sangon Biotech, Bloom 238∼282 g, Shanghai), Type-B Galetin (A600908-0500, Sangon Biotech, Shanghai), Type-A Galetin (S25197-100g, Yuanye Bio-Technology, Shanghai), Type-B Galetin (S22176-100g, Bloom ∼223 g, Yuanye Bio-Technology, Shanghai), Type-A Galetin (V900863-100G, Bloom ∼300 g, Sigma), Type-B Galetin (G9391-100G, Bloom ∼225 g, Sigma), 88% Formic acid (10010118, SinoPharm, Beijing).

### Bone sample collection

Male C57BL/6J mice (4 weeks to 8 months old) were obtained from Jiangsu GemPharmatech Co., Ltd. (Jiangsu, China) and housed under standard conditions. A mouse bone growth retardation model was established as described in our previous studies [13, 15]. Femurs and tibiae from mice were collected after euthanasia via carbon dioxide inhalation. Femurs from rabbit (*Oryctolagus cuniculus*), pig (*Sus scrofa domesticus*), beef cattle (*Bos taurus domesticus*), bullfrogs (*Lithobates catesbeiana*), chicken (*Gallus gallus domesticus*), and wall lizard (*Gekko* sp.), as well as caudal vertebrae from common carp (*Cyprinus carpio*) and little yellow croaker (*Larimichthys polyactis*), were isolated upon purchase. All animal experiments complied with institutional ethical guidelines and were approved by the Animal Experiments Ethical Committee of Shanghai Municipal Hospital of Traditional Chinese Medicine (Approval No. 2023107).

### Bone sample processing

After fixation in 10% neutral formalin at room temperature for 24 hours, femur, tibia, and vertebral bone samples were decalcified in various decalcification solutions (10% EDTA, 14% EDTA, 0.5 mol/L EDTA, and 5% formic acid) at room temperature for 7, 14, 21, and 28 days on a shaker, with the solution replaced daily within the 7-day decalcification period, or every 3 days for decalcification periods beyond 7 days. Following decalcification, the bones were dehydrated, paraffin–embedded, and sectioned at 4 μm thickness.

### OLCS silver nitrate staining

The OLCS silver nitrate staining was performed according to our previously reported methods [13–15]. Briefly, 1 mol/L silver nitrate was mixed with 2% (w/v) type-B gelatin in 1% (v/v) formic acid at a 2:1 (v/v) ratio. Unless otherwise specified in Figure 5, Solarbio™ type-B gelatin was used for all experiments in this study. The mixture was immediately applied to bone paraffin sections, which were then placed on the lid of a high-precision water bath (Model DKB-501S, Jinghong, Shanghai) filled with tap water, or alternatively on a constant-temperature hotplate or in a constant–temperature incubator. Staining was performed at various temperatures (37°C, 40°C, 45°C, 50°C, 55°C, 60°C, 65°C, and 70°C) for 5, 10, or 20 minutes. Two control conditions were employed: (1) room temperature staining under 254 nm ultraviolet (UV) irradiation (Biological Safety Cabinet, 1379/1389 with 254 nm UV light G3675L 31–40 W, Thermo) for 60 minutes, and (2) the Ploton silver method (50% silver nitrate solution) with room temperature staining in the dark for 55 minutes. After the staining steps, sections were rinsed directly with 18.2 MΩ·cm Milli-Q water to stop the staining reaction, without the use of 5% sodium thiosulfate.

### OLCS Quantification

OLCS images were captured at 400x magnification using a Nikon bright-field microscope (ECLIPSE 200, Nikon) and quantified using ImageJ 1.53k software (NIH). The quantification process was conducted as previously reported in our research [13–15] as follows: first, the image was processed by selecting *Image > Color > Split Channels > Green* to isolate the green channel. Next, contrast enhancement was performed by navigating to *Process > Enhance Contrast*, setting the saturated pixels to 0.3%, and equalizing the histogram. The threshold was then adjusted using *Image > Adjust > Threshold*, followed by applying the changes. The region of interest (ROI) was defined using the Polygon selection tool, and measurement parameters were set by selecting *Analyze > Set Measurements*, which included *Area*, *Integrated Density*, *Area Fraction*, *Mean Gray Value*, and *Limit to Threshold*. Finally, the positive staining area was measured as a percentage of the LCS area relative to the bone area (*% area*) using *Analyze > Measure*. This procedure was repeated for all images.

## Data reproducibility and statistics analysis

For all qualitative comparisons presented in Figures 1–5, each staining condition was replicated on at least two bone paraffin sections, with one to two sections per slide. For the quantitative analyses shown in Figure 6, each staining condition was replicated on six slides, including one to two bone paraffin sections per slide. From the cortical bone region of each longitudinal femoral or tibial section, a minimum of 3–5 random regions of interest (ROIs) were captured at 400× magnification. The quantified data of the ratio of LCS area to bone area (%) obtained from ImageJ were imported into GraphPad Prism 8.0.1 software for quantitative analysis. Normality was assessed with the Shapiro–Wilk test, and group comparisons were performed using one-way ANOVA followed by Tukey’s multiple comparisons. Data are presented as means ± standard deviation (SD), with error bars representing the SD. Statistical significance was defined as *p* < 0.05.

**Figure 1.**
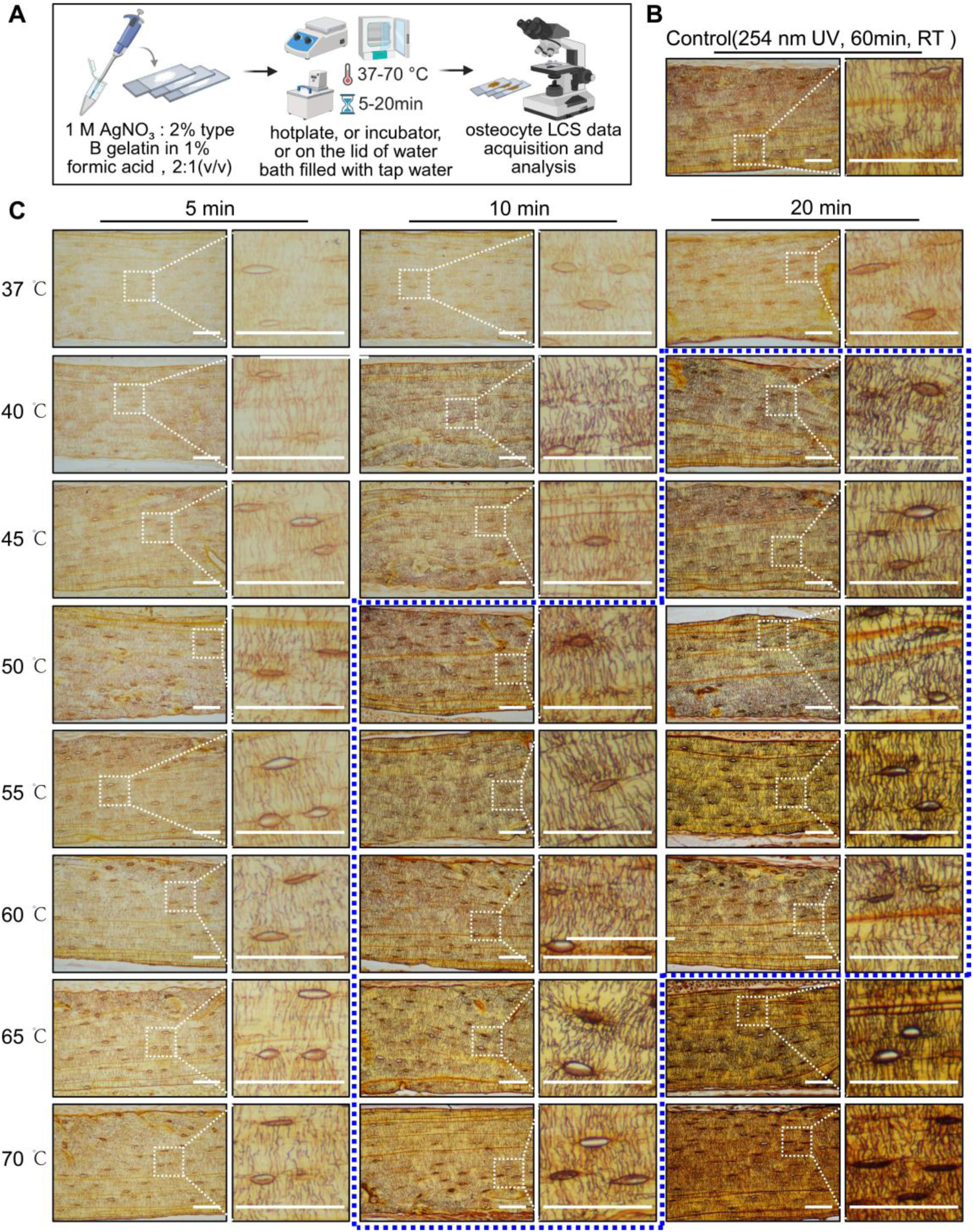
Effects of staining temperature on silver nitrate staining of the osteocyte LCS. (A) LCS staining workflow, created with BioRender (https://BioRender.com). (B) LCS staining at 254 nm UV irradiation for 60 minutes. All scale bars = 50 μm. (C) LCS staining at different temperatures for various durations. All scale bars = 50 μm. The dashed blue box indicates the optimal staining temperature and corresponding duration for effective staining of osteocyte LCS.

**Figure 2.**
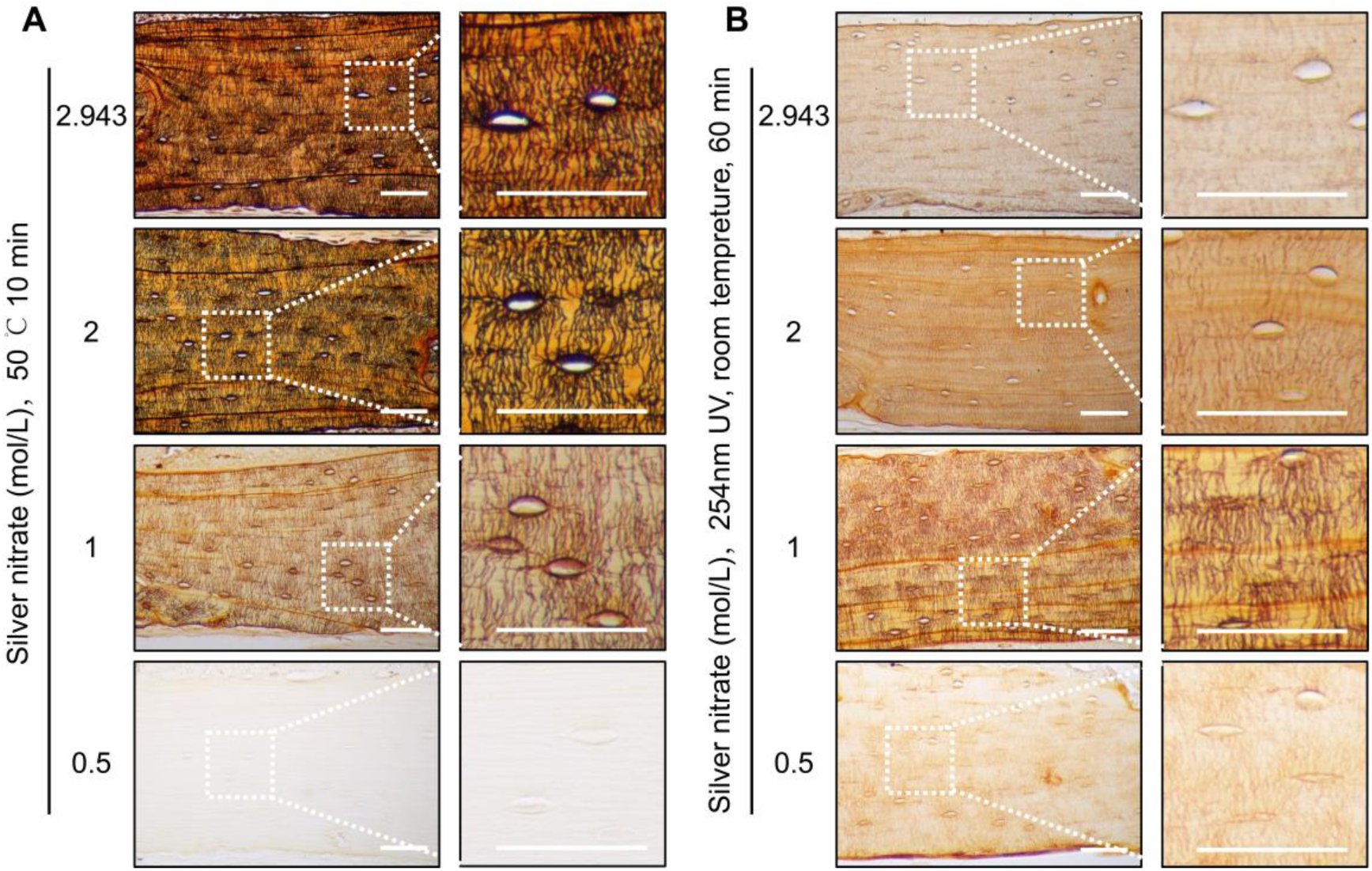
Comparison of LCS staining performance using different concentrations of silver nitrate at 50 °C for 10 minutes. (A) LCS staining performance with different concentrations of silver nitrate at 50 °C for 10 minutes. All scale bars = 50 μm. (B) LCS staining performance with different concentrations of silver nitrate at room temperature under 254 nm UV irradiation for 60 minutes. All scale bars = 50 μm.

**Figure 3.**
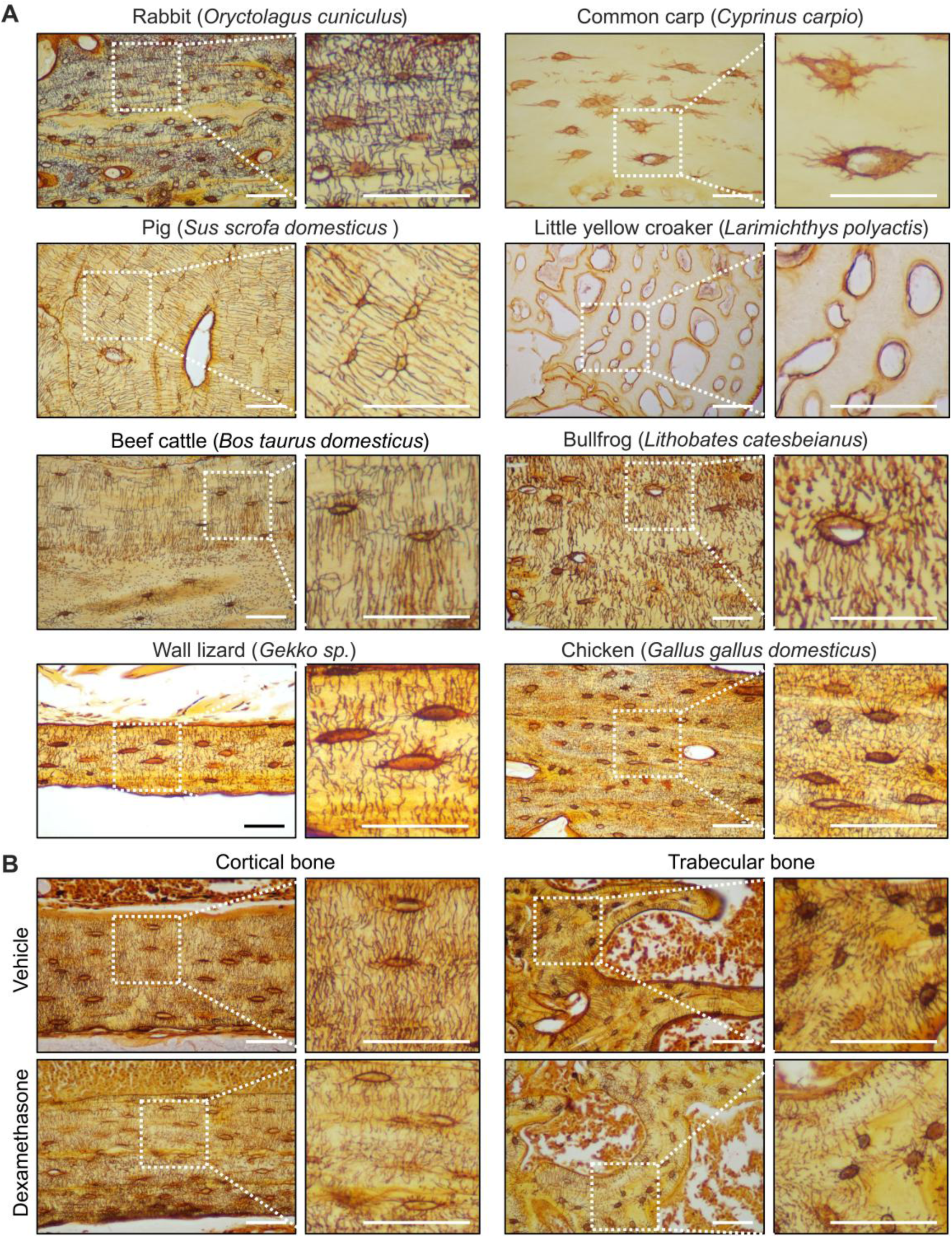
LCS staining performance using 1 mol/L silver nitrate at 50 °C for 10 minutes in bones from different vertebrate species and a mouse pathological bone model. (A) LCS staining performance in bones from various vertebrate species. All scale bars = 50 μm. (B) LCS staining performance in cortical and trabecular bone of the GC-treated mice. All scale bars = 50 μm.

**Figure 4.**
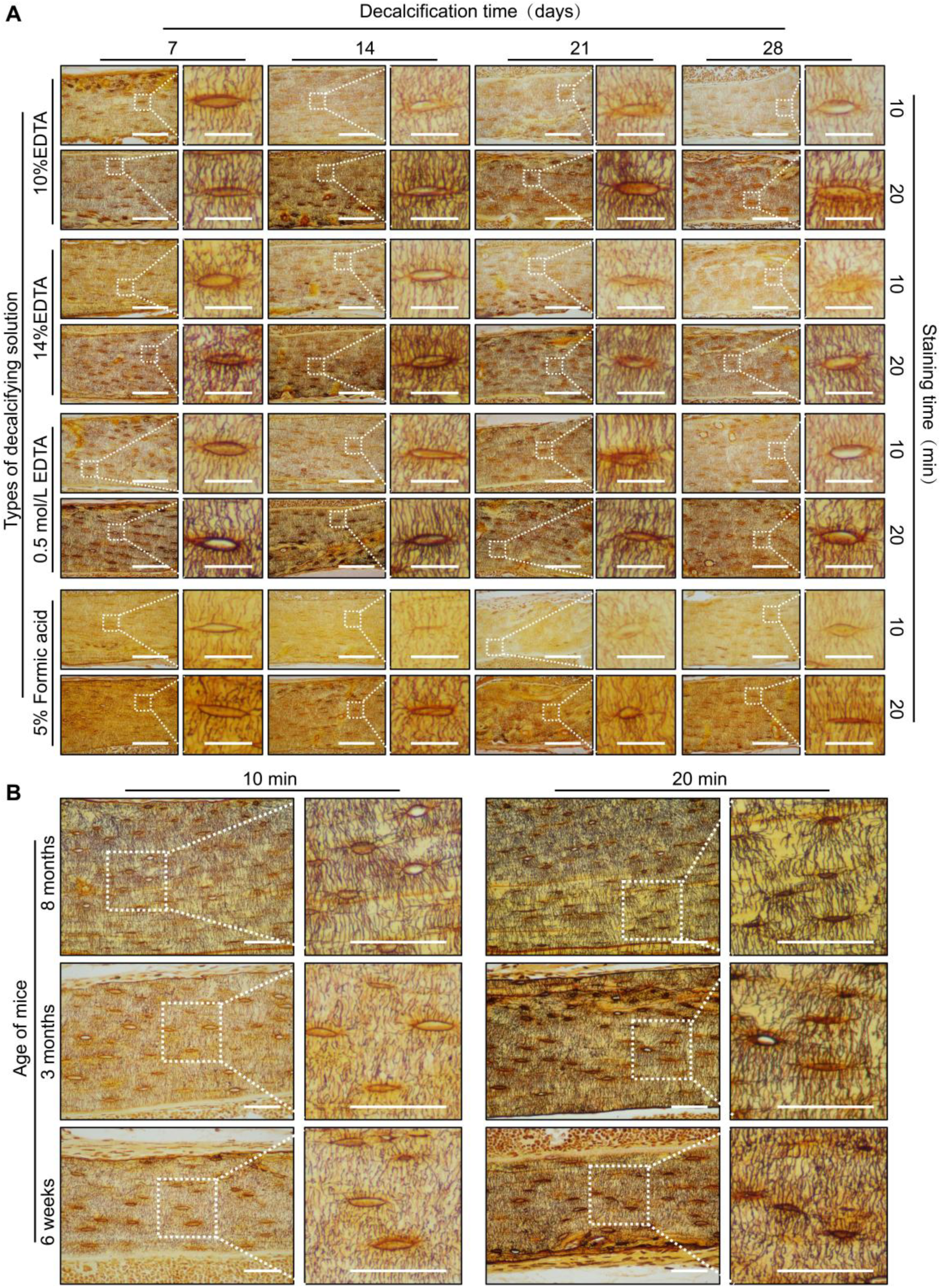
LCS staining performance using 1 mol/L silver nitrate at 50 °C for 10–20 minutes on bone tissues across different ages and various decalcifying solutions. (A) LCS staining performance of bones decalcified using different decalcifying solutions. Scale bars = 100 μm (large images) and 20 μm (insets). (B) LCS staining performance of bones from mice of different ages. All scale bars = 50 μm.

**Figure 5.**
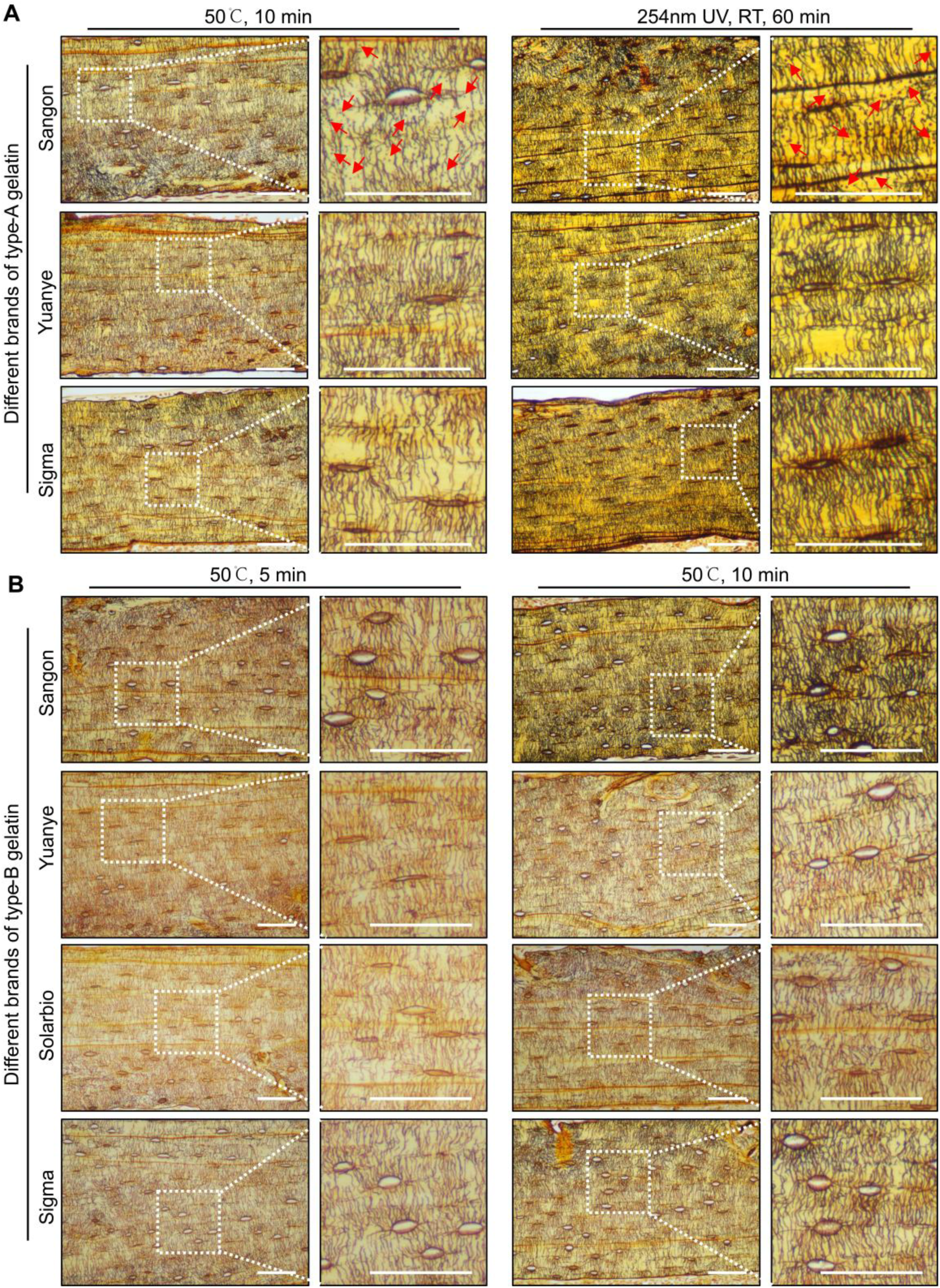
Effect of different gelatin types and commercial sources on LCS staining performance of the rapid Wu–Wang Silver Method. (A) LCS staining performance achieved using type-A gelatin from different commercial manufacturers. Red arrows indicate black granular precipitates. All scale bars = 50 μm. (B) LCS staining performance achieved using type-B gelatin from different commercial manufacturers. All scale bars = 50 μm.

**Figure 6.**
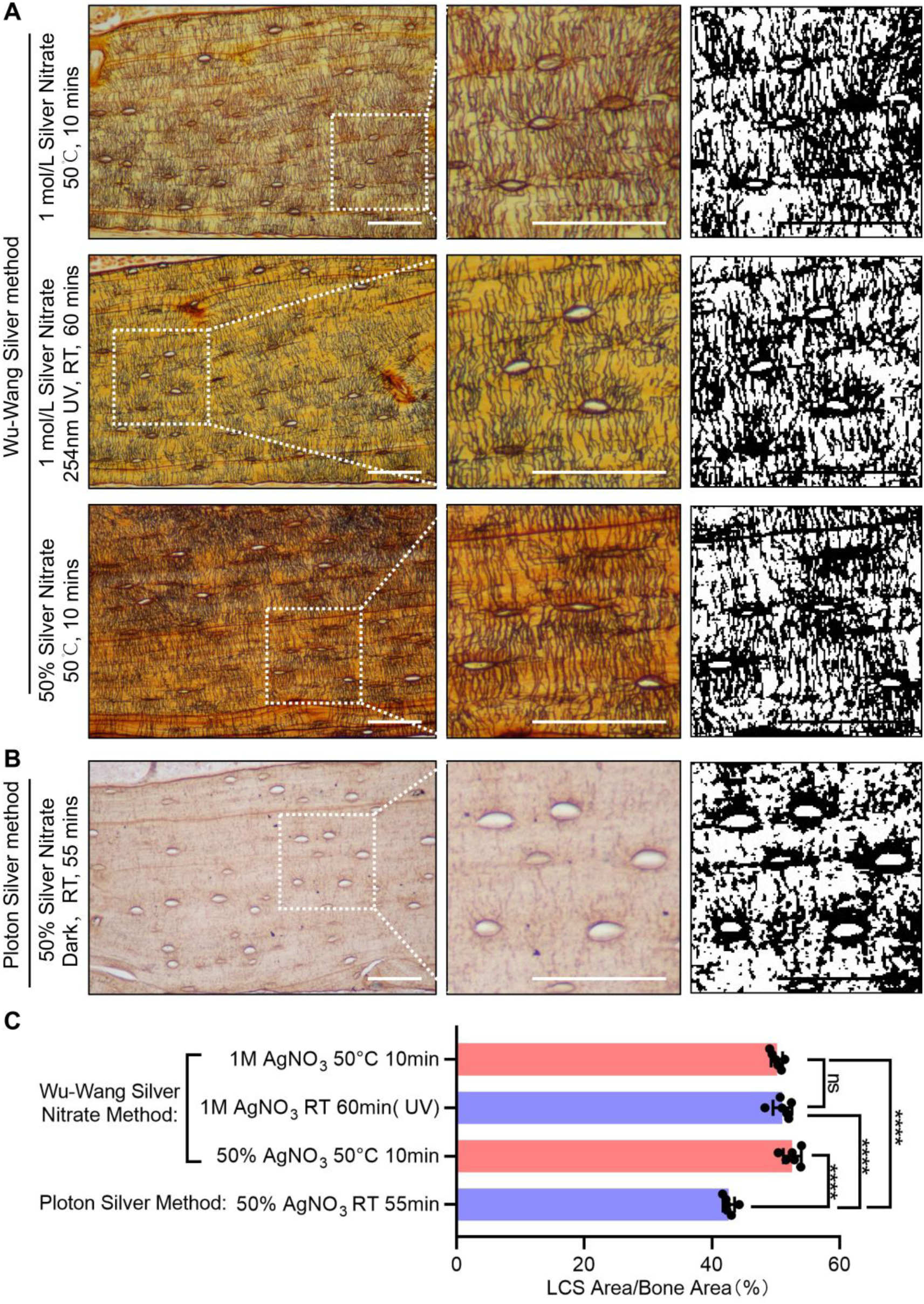
Comparison of the Wu–Wang silver method and the Ploton silver method for osteocyte LCS visualization. (A) Staining performance of the rapid Wu–Wang silver method. All scale bars = 50 μm. (B) Staining performance of the classic Ploton silver method. All scale bars = 50 μm. (C) Quantitative analysis of LCS density between two methods, analyzed by one-way ANOVA. Data are presented as mean ± SD (n = 6). ****p < 0.0001, ns = not significant.

## Results

### 1. Incubation at 50–70 °C for 10 minutes enables efficient staining of the osteocyte LCS

In our previous studies, we observed that elevating the staining temperature could shorten the required duration [15]. We therefore hypothesized that a systematic comparison of the effects of different temperatures on LCS staining could yield a more efficient method than our previously optimized protocol, while simultaneously overcoming the inherent limitations of the Ploton silver method and further advancing osteocyte and LCS research. Based on this idea, we systematically evaluated how various staining temperatures influence the outcomes of LCS silver nitrate staining. Staining was performed on femurs and tibiae harvested from 8-month-old mice and decalcified in 0.5 M EDTA for 7 days, using a 1 mol/L silver nitrate solution (Figure 1A). As a control, parallel staining was carried out at room temperature under 254 nm UV irradiation for 60 minutes (Figure 1B).

Our results showed that elevating the staining temperature from room temperature to 37–70 °C enabled discernible brown LCS staining in the cortical bone region after only 5 minutes of incubation. Moreover, both staining intensity and clarity increased significantly as the temperature rose from 37 °C to 70 °C (Figure 1C). At the same temperatures, extending the incubation time to 10 or 20 minutes further enhanced staining intensity and clarity. Notably, incubation for 10 minutes at 50–70 °C (Figure 1C) or for 20 minutes at 40–60 °C (Figure 1C) produced staining intensity and clarity comparable to those obtained with our previously optimized method (Figure 1B).

However, when sections were stained at 65–70 °C for 20 minutes, the samples became overly dark and the LCS signal appeared subdued. Many lacunae and the surrounding peri-lacunar regions were filled with brown–black precipitates, consistent with an overall over-staining effect (Figure 1C). At 80 °C, severe tissue detachment from the slides and rapid evaporation of the staining solution were observed (data not shown), indicating that excessively high temperatures are unsuitable for LCS staining.

Overall, the qualitative comparisons indicate that elevated staining temperatures substantially reduce the time needed for LCS silver nitrate staining—from 55–60 minutes to under 10 minutes. Specifically, incubation with 1 mol/L silver nitrate at 50–70 °C for 10 minutes or at 40–60 °C for 20 minutes achieves optimal staining (dashed blue box in Figure 1C).

### 2. 1–2.943 mol/L silver nitrate incubation at 50 °C for 10 minutes enables efficient LCS staining and mitigates the Ploton silver method–associated LCS destruction and insufficient staining

After identifying staining temperature as the key factor for improving staining efficiency and defining an optimal temperature–time window, we next compared the staining performance of different silver nitrate concentrations under high-temperature conditions. Staining at 50 °C for 10 minutes was chosen as the standard condition. Staining was performed on femurs and tibiae harvested from 8-month-old mice and decalcified in 0.5 M EDTA for 7 days using silver nitrate solutions at concentrations of 0.5, 1, 2, and 2.943 mol/L (equivalent to 50% w/v silver nitrate as used in the Ploton silver staining method) (Figure 2A). As a control, parallel staining was carried out at room temperature under 254 nm UV irradiation for 60 minutes (Figure 2B).

The results showed that, relative to staining under 254 nm UV irradiation for 60 minutes (Figure 2B), 1, 2, and 2.943 mol/L silver nitrate solutions all yielded clear LCS staining at 50 °C for 10 minutes (Figure 2A). Moreover, higher silver nitrate concentrations produced stronger staining intensity, but also resulted in elevated background signal (Figure 2A). The staining quality achieved with 1 mol/L silver nitrate at 50 °C for 10 minutes was comparable to that obtained under 254 nm UV irradiation for 60 minutes (Figure 2A). In contrast, 0.5 mol/L silver nitrate at 50 °C for 10 minutes produced almost no effective LCS staining (Figure 2A). Notably, staining at 50 °C for 10 minutes unexpectedly overcame the insufficient or destructive LCS staining observed with high-concentration silver nitrate (2 and 2.943 mol/L) under room-temperature 254 nm UV irradiation for 60 minutes (Figure 2A).

Therefore, silver nitrate solutions within the range of 1–2.943 mol/L reliably produce clear and effective LCS staining at 50 °C for 10 minutes. Furthermore, high-temperature staining circumvents the inherent limitations of high-concentration silver nitrate solutions during prolonged room-temperature staining.

### 3. 1 mol/L silver nitrate incubation at 50 °C for 10 minutes enables efficient LCS staining in bones from diverse vertebrate species and mouse pathological models

To assess the broad applicability of our rapid silver staining method, we evaluated its performance on bone tissues from multiple vertebrate species and on mouse pathological models. Staining at 50 °C for 10 minutes using a 1 mol/L silver nitrate solution was selected as the standard condition. Staining was performed on femur, tibia, and vertebral bone samples harvested from representative vertebrate taxa and from a mouse model of glucocorticoid (GC)-induced bone growth retardation. All samples were decalcified in 0.5 mol/L EDTA for 7 days (Figure 3).

Our results showed that incubation with 1 mol/L silver nitrate at 50 °C for 10 minutes effectively stained the LCS in mammals (e.g., rabbit, pig, beef cattle), amphibians (e.g., bullfrog), birds (e.g., chicken), reptiles (e.g., wall lizard), and fishes (e.g., common carp, little yellow croaker) (Figure 3A). In accordance with our previous findings, vertebrae of the little yellow croaker (*Larimichthys polyactis*) lacked osteocytes and elaborate LCS structures, presenting only cavity-like features, in contrast to the common carp (*Cyprinus carpio*). Furthermore, this rapid staining method also enabled efficient LCS staining in bones from a mouse model of GC-induced bone growth retardation (Figure 3B). Compared with control mice, LCS was markedly reduced in both cortical and trabecular bone of the GC-treated mice, which was consistent with our previous results [13, 15].

Therefore, 1 mol/L silver nitrate at 50 °C for 10 minutes provides efficient LCS staining for bone tissues across diverse vertebrate species and common mouse bone pathological models.

### 4. 1 mol/L silver nitrate at 50 °C for 10–20 minutes enables effective LCS staining of bone tissues across different ages and various decalcifying solutions

We first established this rapid and effective staining method in paraffin sections of mouse long bones (femurs and tibiae) from 8-month-old animals after decalcification for 7 days in 0.5 mol/L EDTA (Figure 1–2). We then considered that mineralization levels vary across growth stages and that different decalcifying agents exhibit distinct decalcification rates [21–22]. Therefore, we anticipated potential staining variability among samples. Therefore, we anticipated potential staining variability across samples. To evaluate this, we assessed staining performance using four commonly used decalcification solutions—10% EDTA, 14% EDTA, 0.5 mol/L EDTA, and 5% formic acid—followed by decalcification for 7, 14, 21, or 28 days in femurs and tibiae from 3-month-old mice. In parallel, femurs and tibiae from mice at 6 weeks, 3 months, and 8 months of age were decalcified for 7 days in 0.5 mol/L EDTA. Staining was performed using 1 mol/L silver nitrate, incubated at 50°C for 10 or 20 minutes (Figure 5).

The results showed that when staining was performed at 50°C for 10 minutes, the staining quality of the osteocyte LCS declined progressively with increasing decalcification time, accompanied by a marked reduction in staining contrast (Figure 5A). When the bones were decalcified for only 7 days using the three EDTA-based solutions, LCS staining was already clear and effective after 10 minutes at 50°C (Figure 5A). When the staining time was extended to 20 minutes, all specimens decalcified with EDTA solutions exhibited clear and effective LCS staining, except those treated with 5% formic acid (Figure 5A). For samples decalcified with 5% formic acid, although staining for 20 minutes was better than that for 10 minutes, the LCS staining remained less clear than in EDTA-decalcified samples and fewer LCS signals were observed (Figure 5A). In addition, compared with 6-week-old and 3-month-old mice, clear and effective LCS staining could be achieved in 8-month-old mice after staining at 50°C for 10 minutes. When the staining time was extended to 20 minutes, clear and effective staining was achieved across all these samples (Figure 5B).

Therefore, the quality of rapid silver staining varies across bone samples from mice of different ages and with different decalcification treatments. However, extending the staining time to 15–20 minutes results in consistent, high-quality LCS staining in all specimens.

### 5. Effect of gelatin type and commercial source on LCS staining performance with 1 mol/L silver nitrate at 50°C for 10 minutes

Our previous studies demonstrated that different gelatin types significantly impact osteocyte LCS staining quality: type-A gelatin frequently induced black granular precipitates in bone sections, a defect not observed with type-B gelatin [13, 15]. Given that the 50°C staining condition not only substantially shortens the required incubation time to 10 minutes, but also prevents the LCS structural damage and insufficient staining seen with high-concentration silver nitrate solutions at room temperature (Figure 2), we sought to determine whether high-temperature staining could mitigate the inherent staining defects of type-A gelatin. Furthermore, we investigated whether LCS staining outcomes were consistent across different commercial sources of both type-A and type-B gelatin. To address these questions, we systematically compared the LCS staining efficacy of type-A gelatin from three commercial manufacturers (Sangon^TM^, Yuanye ^TM^, Sigma ^TM^) and type-B gelatin from four manufacturers (Sangon^TM^, Solarbio^TM^, Yuanye ^TM^, Sigma ^TM^) (Figure 5). Staining was performed on femurs and tibiae harvested from 8-month-old mice, which were decalcified in 0.5 mol/L EDTA for 7 days. Samples were stained with a 1 mol/L silver nitrate solution, incubated at 50°C for 5 or 10 minutes.

A considerable variation in LCS staining outcomes was observed among the three type-A gelatin brands (Figure 5A). Consistent with our previous reports [13,15], type-A gelatin from Sangon^TM^ produced abundant black granular precipitates in stained sections, regardless of whether incubation was performed at room temperature under 254 nm UV irradiation for 60 minutes, or at 50°C for 10 minutes (Figure 5A). In contrast, type-A gelatin from Yuanye ^TM^ and Sigma ^TM^ did not generate such precipitates under either incubation condition (Figure 5A). Among the four type-B gelatin manufacturers tested, Sangon^TM^ type-B gelatin achieved equivalent LCS staining efficacy in only 5 minutes, whereas the other three brands typically required 10 minutes (Figure 5B). When incubation time was extended to 10 minutes, Sangon™ type-B gelatin consistently produced clearer LCS staining than the other three type-B gelatins (Figure 5B).

These results indicate that high-temperature staining does not circumvent the inherent defect associated with specific brands of type-A gelatin (e.g., Sangon™). However, selecting an appropriate type-A gelatin brand (e.g., Yuanye™ or Sigma™) can prevent the formation of black granular precipitates under both high-temperature and room-temperature staining conditions. In addition, LCS staining efficiency at elevated temperatures varies among different commercial type-B gelatin products. Our results show that using an appropriate type-B gelatin brand (e.g., Sangon™), enables clear and effective rapid LCS staining within 5 minutes.

### 6. The Wu–Wang silver method provides superior LCS visualization compared with the Ploton silver method

We next quantitatively compared the number of stained LCS structures produced by the high–temperature Wu–Wang silver method and the room–temperature Ploton silver method. Staining was performed on femurs and tibiae from 8-month-old mice that were decalcified in 0.5 M EDTA for 7 days. Samples were stained using either 1 mol/L silver nitrate or 50% (w/v) silver nitrate (2.943 mol/L) at 50°C for 10 minutes or under 254 nm UV irradiation at room temperature for 60 minutes (Figure 6A). For comparison, the Ploton silver method (55 minutes at room temperature in the dark) was used as the control (Figure 6B).

The results showed that 1 mol/L silver nitrate at 50°C for 10 minutes produced efficient LCS staining comparable to that obtained with 1 mol/L silver nitrate under 254 nm UV irradiation at room temperature (RT) for 60 minutes (Figure 6A). The LCS staining quality achieved under both conditions was superior to that obtained with the classic Ploton silver staining method (Figure 6B). Meanwhile, 50% (w/v) silver nitrate at 50°C for 10 minutes also yielded markedly improved LCS staining outcomes compared with the Ploton silver method performed at RT for 55 minutes in the dark (Figure 6B). As we previously reported, the Ploton silver method either fails to stain the LCS efficiently or causes structural disruption of the LCS, resulting in an intermittent, irregular, and blurred LCS pattern in cortical bone (Figure 6B). Quantification of LCS density further showed that staining with 1 mol/L silver nitrate at 50°C for 10 minutes produced LCS densities comparable to those obtained with 1 mol/L silver nitrate under 254 nm UV irradiation at RT for 60 minutes. Furthermore, all three optimized staining conditions yielded significantly higher LCS density than the classic Ploton silver method performed at RT for 55 minutes in the dark.

Taken together, these results indicate that our high-temperature rapid Wu–Wang silver method significantly outperforms the Ploton silver method, and overcomes the key limitations of Ploton silver method, including insufficient staining and potential structural disruption of the LCS.

## Discussion

In this study, we report a novel rapid silver nitrate staining method that enables efficient staining of the osteocyte LCS in vertebrate skeletal tissues within 5-10 minutes at 50–70 °C. This rapid method is markedly superior to the Ploton silver method, and effectively overcomes its inherent limitations, including incomplete staining and structural damage to the LCS.

The silver nitrate staining method was originally developed to visualize chromosomal nucleolar organizer regions (NORs) [23]. This structure was first described by Heitz in 1931 [24] and subsequently by McClintock in 1934 [25]. Following silver staining, NORs are referred to as argyrophilic nucleolus organizer regions (AgNORs) [23]. In 1975, Goodpasture and Bloom introduced a specific staining method for NORs using 50% silver nitrate, ammoniacal silver solution, and a 3% formalin developing solution [26–27]. In their protocol, after application of 50% silver nitrate to slides, the slides were placed 25 cm below a photo flood lamp (2800 K bulb) for 10 minutes. Since a 2800 K bulb produces substantial radiant heat [28], we infer that the actual staining temperature during this step would have been substantially higher than room temperature. Building on the 50% silver nitrate formulation from Goodpasture and Bloom, Howell and Black developed a one-step silver staining method in 1980, with a formulation (50% w/v silver nitrate, 2% w/v gelatin in 1% v/v formic acid) that remains widely used today [29]. In this method, staining was performed at 70°C for 2 minutes. Following staining, the authors noted that “the slide background is homogeneously clean, with little or no extraneous silver precipitate” [29].

In 1986, Ploton et al. were the first to adapt the Howell and Black silver staining method to tumor pathology, and demonstrated that the number of interphase AgNORs in human prostate cancer cells was significantly higher than that in corresponding benign or hyperplastic cells [23, 30]. In this pioneering work, Ploton et al. noted that when staining was performed at 60°C, “slight nonspecific staining was noticeable; this gave the slide a general brownish colour visible to the naked eye” [30]. They further attributed this background staining to “nonspecific adsorption and/or precipitation of silver due to the high temperature of the reaction” [30]. To overcome this issue, Ploton et al. reduced the staining temperature to 20°C. Under these conditions, the authors observed that the required staining duration was prolonged, which in turn made the reaction easier to monitor in real time. Critically, they reported no background staining for any of the tested materials, with staining signal strictly confined to nucleolar regions and mitotic NORs [30]. In 1996, Chappard et al. applied Ploton’s room-temperature staining conditions to NOR staining in bone sections, and unexpectedly found that this method achieved excellent staining of the osteocyte LCS [11–12]. In this study, Chappard et al. established the staining conditions that have been widely used in osteocyte LCS research ever since: “55 minutes at room temperature in the dark” [11–12].

In our previous report, we observed that increasing the staining temperature may accelerate the silver staining reaction [15]. Building on this observation and established evidence concerning the temperature dependence of silver impregnation [31–33], we systematically evaluated staining performance for the osteocyte LCS under different temperature conditions. We found that raising the staining temperature substantially shortened staining time from the conventional 60 minutes (or 55 minutes) to within 10 minutes. With 1 mol/L silver nitrate, clear and effective LCS staining (as indicated by the dashed blue box in Figure 1A) was achieved either by staining for 10 minutes at 50–70°C or by staining for 20 minutes at 40–60°C. However, when the temperature was further increased (e.g., 80°C), bone sections exhibited severe detachment from slides, and the staining solution underwent rapid evaporation, indicating that excessively high temperatures are not suitable for LCS staining. Comparisons across different silver nitrate concentrations demonstrated that solution concentrations ≥ 1 mol/L produced effective LCS staining after 10 minutes at 50°C. Notably, and contrary to our expectations, incubation at 50°C for 10 minutes almost fully mitigated the LCS damage and insufficient staining caused by high silver nitrate concentrations (2 mol/L and 2.943 mol/L) under room-temperature incubation conditions (Figure 2). Overall, staining with 1 mol/L silver nitrate at 50°C–70°C for 10 minutes provides a rapid and robust method for achieving clear and effective LCS visualization.

Next, we applied this novel rapid staining method to paraffin-embedded bone sections from diverse vertebrate species and mouse models of bone diseases. Our data further showed that incubation with 1 mol/L silver nitrate for 10 minutes at 50°C enables clear and effective LCS staining of both cortical and trabecular bone in a glucocorticoid (GC)-induced mouse model of bone growth retardation. Remarkably, this rapid method also enables clear and effective LCS staining across all five major vertebrate taxa: mammals, birds, reptiles, amphibians, and fish. Consistent with our previous observations [13, 15], fish exhibit strikingly distinct skeletal architecture. Compared with the dense and complex osteocyte network and LCS found in tetrapod mammals, fish such as the common carp (*Cyprinus carpio*) typically contain far fewer osteocytes and simpler LCS structures (Figure 3A). Furthermore, osteocytes and the LCS show striking contrasts among teleosts of different taxonomic groups. Compared with basal teleosts (e.g., common carp, *Cyprinus carpio*), vertebrae of advanced teleosts (little yellow croaker, *Larimichthys polyactis*) lack osteocytes and a complex LCS, exhibiting only cavity-like structures (Figure 3A). Together, these findings address a fundamental question: what is the biological significance of the apparent lack of osteocytes and LCS in advanced teleosts [16–18]? In particular, why do some teleosts retain osteocytes and LCS, whereas others display severe reduction or complete absence of these structures [16–18]? Alternatively, from an evolutionary perspective, how did the novel osteocyte cell type and its LCS emerge across tetrapod lineages—amphibians, reptiles, terrestrial mammals, and birds—which all evolved from teleost ancestors [19–20]? Addressing these questions through comparative studies may provide critical insight into how animals evolve at the cellular level to adapt to distinct living environments. It may also help clarify a broader evolutionary problem: the origin, loss, and re-emergence of specific cell types and their associated microanatomical structures [19–20].

In addition, we performed a systematic comparison of variables that may influence the performance of our rapid staining method, including decalcification conditions, bone sample age, and gelatin brands/manufacturers. To directly assess the effect of decalcification, we used four different decalcification solutions to decalcify femurs and tibiae from 3-month-old mice for different durations (7, 14, 21, and 28 days). We found that when sections were stained at 50°C for 10 minutes, the staining quality of the osteocyte LCS declined progressively with increasing decalcification time, accompanied by a marked reduction in staining contrast. When staining time was extended to 20 minutes, all EDTA-decalcified samples produced clear and effective LCS staining, except for samples decalcified using 5% formic acid. Notably, 5% formic acid, despite being widely used as a rapid decalcification solution, was found to cause LCS structural disruption, making it poorly suited for LCS silver staining. Therefore, for LCS analysis, we recommend using an EDTA-based decalcification solution. Decalcification for 7 days was sufficient to obtain well-prepared sections that could be efficiently and rapidly stained within 10 minutes using our rapid method. When the decalcification duration was extended beyond 7 days, we recommend increasing the silver-staining time to 15–20 minutes to achieve efficient LCS visualization.

A qualitative comparison of the performance of our rapid silver staining method for LCS visualization in paraffin sections from 6-week-, 3-month-, and 8-month-old mice revealed an age-dependent difference in staining efficacy. At 50°C for 10 minutes, the 8-month-old group showed more distinct and robust LCS staining than the 6-week- and 3-month-old groups. When the staining time was extended to 20 minutes, LCS staining efficacy in the 6-week- and 3-month-old groups became comparable to that of the 8-month-old group. We hypothesize that this modest discrepancy reflects age-related differences in bone mineralization [21]. In general, after decalcification under identical conditions, younger bones retain fewer residual mineral salts than more mature bones [22], which could subtly affect the efficiency of our rapid LCS silver staining method. Overall, for LCS analyses across developmental stages, we recommend extending the staining time to 15–20 minutes to ensure clear and reliable visualization.

A key contribution of the method developed by Howell and Black was the introduction of gelatin into the silver staining system [29]. In their method, 2% (w/v) gelatin combined with 1% (v/v) formic acid was formulated as a protective colloidal developer. This solution helps to regulate the staining reaction and, importantly, reduces background staining that arises from the intense reduction of silver nitrate [29]. In our previous study, we found that the osteocyte LCS could be stained when 0.05–10% (v/v) formic acid was used alone (without gelatin) in combination with 1 mol/L silver nitrate. However, bone sections exhibited abundant black granular precipitates adhering to the slides, which rendered the data unusable [13, 15]. By contrast, when 0.05–2% type-B gelatin was included in the formic acid, the LCS could be stained clearly and effectively, with no black granular precipitates. These observations indicate that gelatin plays a critical role in the silver nitrate staining method. As Howell and Black noted in their original paper: “until now, however, cytogeneticists have not used protective colloids to control silver-staining of Ag-NORs.” They were the first to apply such gelatin-based protective colloids to the silver staining of NORs. Owing to this pivotal improvement, silver staining has since been widely adopted by cell biologists and histologists, and is widely regarded as “the simplest and most replicable technique” [34].

In our previous study, we demonstrated that type-B gelatin enables effective LCS staining [13, 15]. Although type-A gelatin can also yield effective LCS staining, bone sections exhibited numerous black granular precipitates that rendered the data unusable. In this study, we unexpectedly found that staining at 50 °C for 10 minutes completely overcame the LCS damage and insufficient staining observed when high concentrations of silver nitrate (2 mol/L and 2.943 mol/L) were applied at room temperature for 60 minutes (Figure 2A). This raised an important question: can elevated-temperature staining also rescue the staining defects associated with type-A gelatin? Moreover, do different brands/manufacturers of type-A and type-B gelatin produce distinct staining outcomes? Our data showed that high-temperature staining did not prevent the formation of black granular precipitates when using certain type-A gelatin products, specifically Sangon™. In contrast, type-A gelatin from Yuanye™ and Sigma™ allowed effective LCS staining without black granular precipitates under both high-temperature and room-temperature conditions. We also observed brands/manufacturers-depende differences in the performance of type-B gelatin. For type-B gelatin from Sangon™, efficient and clear LCS staining was achieved in as little as 5 minutes at 50 °C.

To our knowledge, this is the first report demonstrating such strong brand- and manufacturer-dependent differences in gelatin performance with respect to LCS silver staining quality and efficiency. Accordingly, by selecting an appropriate brand of type-B gelatin (e.g., Sangon™), we were able to substantially reduce the required staining time—from 60 minutes or 55 minutes to within just 5 minutes. Although type-A gelatin is generally produced by acid hydrolysis of collagen and type-B gelatin by alkaline processing [35–36], our findings indicate that the intrinsic physicochemical properties of gelatin (such as isoelectric point, pH, gel strength, and others), which vary among commercial brands and manufacturers, can directly influence silver staining performance, warranting further investigation. Furthermore, whether there exists a more effective and faster protective colloidal developer than gelatin for staining the osteocyte LCS is also a key question worthy of follow-up research.

Finally, we quantitatively compared the number of LCS structures visualized using our high-temperature staining conditions (the Wu–Wang silver method) versus room-temperature staining conditions (the Ploton silver method). Both qualitative observations and quantitative analyses showed that the LCS densities obtained with the Wu–Wang silver method (50°C for 10 minutes) were comparable to those achieved using 1 mol/L silver nitrate with 254 nm UV irradiation at room temperature for 60 minutes. In parallel, 50% (w/v) silver nitrate at 50°C for 10 minutes produced substantially improved LCS staining compared with the classic Ploton protocol performed at room temperature in the dark for 55 minutes. Notably, all three conditions resulted in significantly higher LCS densities than the Ploton method at room temperature for 55 minutes. Taken together, these findings indicate that the high-temperature Wu–Wang silver method is markedly superior to the Ploton method and effectively overcomes its intrinsic limitations, including LCS structural damage and insufficient staining [13,15]. These findings provide solid support for the widespread acceptance and application of our novel, rapid method by bone biologists investigating the intricate osteocyte LCS.

Although numerous techniques have been developed for staining and imaging the osteocyte LCS, there remains no method capable of generating a whole-bone-level 3D atlas of the osteocyte network and its LCS connectome [10]. Our rapid silver staining method enables effective visualization of submicron-scale LCS structural features using only a standard microscope equipped with 40× or 60× objectives, with resolution comparable to that achieved with the fluorescent dye fluorescein isothiocyanate (FITC) [37] or rhodamine 6G [38] labeling and imaging via laser scanning confocal microscopy (LSCM). However, due to the high density of LCS structures, this method alone cannot fully resolve fine architectural details, including the branching and connectivity of canaliculi. The advent of expansion microscopy (ExM) may provide a practical strategy to overcome this key limitation [39–40]. Specifically, if decalcified bone specimens undergo tissue expansion followed by serial sectioning and staining with the rapid silver nitrate method established in this study, combining this workflow with high-throughput image acquisition, AI-assisted image recognition, and 3D reconstruction could enable the generation of the world’s first complete whole-bone 3D LCS connectome atlas—a landmark goal directly analogous to the ongoing brain connectomics initiatives pursued by neuroscientists worldwide [41].

## Conclusion

In this study, we discovered and established a novel rapid silver nitrate staining method that enables effective staining of osteocyte LCS in bone by incubating in 1 mol/L silver nitrate for 5-10 minutes at 50–70°C. This novel method enables rapid and efficient LCS staining across the skeletons of representative species from all five vertebrate taxa—mammals, birds, reptiles, amphibians, and fish. This novel method, termed the Wu–Wang silver method, stains significantly more LCS than the Ploton silver method. Importantly, it also overcomes LCS structural damage and insufficient staining that can occur with the original Ploton silver method. Given the growing recognition of bone as the body’s largest endocrine organ, and the increasing interest in osteocyte-secreted proteins that mediate remote, long-range regulation of other organs, we expect that broad adoption of this novel, rapid, and efficient method will accelerate research into bone and joint development and disease, as well as other bone metabolic disorders affecting populations worldwide. We further anticipate that this method will attract the interest of evolutionary developmental biologists and paleontologists. By enabling systematic comparisons of LCS morphology, it should facilitate investigations into adaptive differences in the LCS across vertebrate species, thereby deepening our understanding of the morphological evolution and development of osteocytes and LCS.

## Acknowledgments

This research was funded by grants from the National Natural Science Foundation of China (NSFC) (82405480, 82104574), Long-Yi S&T Innovation Cultivation Project of Longhua Hospital Shanghai University of Traditional Chinese Medicine (YD202208), Longhua Hospital Young Talent Training System (XH40204-20250471).

## CRediT authorship contribution statement

Jinlian Wu: Investigation, Validation, Funding acquisition, Formal analysis, Writing-original draft. Libo Wang: Conceptualization, Methodology, Supervision, Funding acquisition, Project administration, Writing-review & editing.

## Dedication

To our parents and daughters, and especially to our mathematics teachers, Mrs. Xiu-Rong Cui and Mrs. Xue-Ying Su: your patience and guidance have taught us that science is a pursuit of beauty and truth through trial and error.

